# ACUTE FOOD DEPRIVATION DECREASES THE EXPRESSION OF METHAMPHETAMINE-INDUCED LOCOMOTOR SENSITIZATION IN RATS

**DOI:** 10.1101/2021.04.20.440661

**Authors:** Juan C Jiménez, Jorge Miranda-Barrientos, Mireya Becerra-Díaz, Florencio Miranda

## Abstract

**INTRODUCTION:** AMPH and METH are known to increase DAergic signaling in the brain reward system by stimulating the release of DA through the reversal of DAT function. However, there is evidence that insulin signaling pathways have the ability to modulate DAT functions. Some studies have reported that hypoinsulinemia attenuates DAT functions, and as a consequence, psychostimulant-induced behaviors are reduced. In the present study, we examined the effects of acute food deprivation, which also reduces insulin levels, on METH-induced locomotor sensitization.

**METHODS:** Separate groups of rats were treated with METH (1 mg/kg i.p.) or saline for 5 days (development phase). On the test day (expression phase), the groups were treated with METH or saline after food deprivation for 24 h. Furthermore, in separate groups of rats, levels of glucose, insulin, and phosphorylation of Akt at Ser473 were also examined after food deprivation for 24 h.

**RESULTS:** The results showed that repeated administration of METH induced a progressive increase in locomotor activity in rats during the development phase. However, METH administration in the expression phase produced a decrease in locomotor activity after 24 h of food deprivation. In addition, the results showed that a reduction in glucose, insulin, and Akt levels occurred as a result of food deprivation.

**CONCLUSION:** These results are in line with previous studies and suggest that food deprivation reduces some behavioral effects of psychostimulants such as AMPH and METH.

**Highlights:** Administration of METH increases DAergic neurotransmission by interfering with proper DA transporter (DAT) function

Some evidence suggests that insulin signaling pathways modulate the DAT in the brain reward system

Acute food deprivation reduces glucose and insulin levels

This study examined the effects of acute food deprivation on METH-induced locomotor sensitization

Present study showed that food deprivation for 24 h reduced METH-induced locomotor sensitization

**Plain Language Summary:** The dopamine transporter (DAT) is responsible for the reuptake of dopamine (DA) and the termination of DA neurotransmission. DAT is also the target for psychostimulant drugs such as cocaine, amphetamine, and methamphetamine. These drugs produce an increase of DA neurotransmission in the reward system. It is now well established that insulin levels modulate DAT function. Several studies have reported that hypoinsulinemia attenuates DAT function, and as a consequence, psychostimulant-induced behaviors are reduced. Food deprivation reduces glucose and insulin levels. In the present study we evaluated the effects of food deprivation on methamphetamine-induced locomotor sensitization. This behavior reflects brain neuroadaptations that contribute to drug addiction. Rats of the main experimental group were treated with methamphetamine for 5 days (development phase). On the test day (expression phase) the rats were treated with methamphetamine after food deprivation for 24 h. In separated groups of rats levels of glucose and insulin were evaluated after food deprivation for 24 h. The results showed that acute food deprivation reduced methamphetamine-induced locomotor sensitization. It was also observed that insulin and glucose levels were reduced after food deprivation. The results of the present study suggest that food deprivation can modulate some behavioral effects of methamphetamine.

## 1. Introduction

The abuse of psychostimulants, such as cocaine, amphetamine (AMPH), and methamphetamine (METH), causes multiple psychiatric disorders including drug addiction. Therefore, it is of fundamental importance to study the neurobiology of psychostimulant-related behaviors. Cocaine, AMPH, and METH are indirect monoamine agonists that exhibit affinity to the dopamine (DA), norepinephrine (NE), and serotonin (5-HT) transporters, which are involved in neurotransmitter reuptake and vesicular storage systems (Proebstl et al., 2019; Rothman & Baumann, 2003). Cocaine is a reuptake inhibitor of DA, NE and 5-HT and therefore increases the synaptic levels of these neurotransmitters. AMPH and METH act on the DA, NE and 5-HT transporters at synaptic vesicles to promote an increase in the cytoplasmic concentration of monoamines and reverse the direction of membrane monoamine transporters, facilitating the efflux of neurotransmitters to the synaptic cleft (Elliot & Beveridge, 2005; Kahlig & Galli, 2003; (Proebstl et al., 2019; Rothman & Baumann, 2003).

The mesolimbic DAergic system, particularly the projection from the ventral tegmental area (VTA) to the nucleus accumbens (nAcc), is an important locus for the production of the locomotor, reinforcing, rewarding, and discriminative stimulus effects of cocaine, AMPH and METH (Di Chiara, 1995; Filip & Cunningham, 2002; Koob, 1992; Pontieri, Tanda, & Di Chiara, 1995). Administration of AMPH, METH or cocaine rapidly increases DAergic neurotransmission by interfering with proper DA transporter (DAT) function and facilitating DAergic signaling in limbic areas (Koob & Bloom, 1988; Koob, 1992; Vaughan & Foster, 2013).

Accumulating evidence suggests that insulin signaling pathways modulate numerous brain functions, including regulation of the DAT in the brain reward system (Daws et al., 2011; Fiory et al, 2019; Jones et al., 2017; Kleinridders & Pothos, 2019; Williams et al., 2007). Some studies have reported the novel ability of insulin signaling pathways in the brain to regulate not only DAT function but also the actions of AMPH (Fiory et al, 2019; Owens et al., 2012). Moreover, some studies have reported that hypoinsulinemia attenuates DAT functions, and as a consequence, psychostimulant-induced behaviors are reduced. For example, Galici et al. (2003) reported that the depletion of insulin by a single injection of streptozotocin reduced AMPH self-administration. In the present study, we examined the effects of acute food deprivation, which also reduces glucose and insulin levels, on the expression of METH-induced locomotor sensitization. Furthermore, the levels of glucose, insulin, and Akt phosphorylation at Ser473 were also examined after acute food deprivation to evaluate these results and reveal the possible mechanism involved in the effects of acute food deprivation on METH-induced locomotor sensitization. Akt is a downstream component of the insulin signaling pathway through activation of PI3K and is involved in the regulation of DA signaling (Beaulieu, Gainetdinov, & Caron, 2007). Akt is reported to play a key role in the surface expression of the DAT (García et al., 2005, Speed et al., 2011), which is required for psychostimulants such as AMPH and METH to enhance DA neurotransmission (Karam & Javitch, 2018; Khoshbouei et al., 2004; McFadden, Vieira-Brock, Hanson, & Fleckenstein 2015). AMPH and its N-methylated derivative METH are psychostimulants that share a common molecular structure (Melega, Williams, Schmitz, Distefano, & Cho, 1995). However, it has been suggested that METH is relatively more potent and addictive than AMPH (Miranda, Cedillo-Ildefonso, Jiménez, Nuñez, & Rodriguez, 2011; NIDA Research Report, 2019; Siefried, Acheson, Lintzeris, & Ezard, 2020). Locomotor sensitization is the progressive and persistent enhancement of a behavioral response to drugs after their repeated and intermittent administration and has been well characterized for psychostimulant drugs (Kalivas & Stewart, 1991; Solis et al., 2021). Furthermore, it is thought to reflect neuroadaptations that contribute to drug addiction and to model some aspects of addictive behaviors such as drug seeking and drug craving (Robinson & Berridge, 1993). Therefore, locomotor sensitization may reflect neurobiological changes related to drug addiction and may be useful for studying DA-insulin interactions. Thus, the main goal of the present study was to examine the effects of acute food deprivation on METH-induced locomotor sensitization in rats.

## 2. Materials and Methods

### 2.1. Animals

Male Wistar rats weighing 220-250 g at the beginning of the experiment were used. The rats were individually housed in standard plastic rodent cages in a colony room maintained at a temperature of 21°C (± 1°C) under a 12 h light/dark cycle (lights on at 6:00 am) and had continuous access to water and food (Teklad LM485 Rat Diet from Harlan, Mexico City, Mexico). All experiments were conducted during the light phase (between 11:00 am and 1:00 pm). Animal care and handling procedures were conducted in accordance with the Official Mexican Norm (NOM-062-ZOO-1999) entitled “Technical Specifications for the Production, Care, and Use of Laboratory Animals”.

### 2.2. Drugs

Methamphetamine hydrochloride (Sigma, St. Louis, MO, USA) was dissolved in saline and injected in a volume of 1 ml/kg via the intraperitoneal (i.p.) route.

### 2.3. Apparatus

Locomotor activity was measured with an open-field activity monitoring system (ENV-515 model; Med Associates, St. Albans, VT, USA). Each Plexiglas cage (40 × 40 × 30 cm) was equipped with 2 sets of 8 photobeams that were placed 2.5 cm above the surface of the floor on opposite walls to record x-y ambulatory movements. Photobeam interruptions were recorded and translated by software to yield the horizontal distance traveled (in cm), which was the dependent measure used for analysis.

### 2.4. Behavioral Procedure

The timeline of the general procedure is shown in Figure 1A. Each day or session started with a 10 min period of habituation to the cages, followed by administration of drug or vehicle (saline). The rats, 10 per group, were returned to the cages, and their locomotor activity was recorded for 60 min. On day −2, the rats were habituated to the open-field cages and injection procedures (habituation session). On day −1, locomotor activity after saline administration (locomotor activity baseline) was evaluated. On days 1-5, the rats in groups M24M and M0M were treated with METH (1.0 mg/kg i.p.) during the development and expression test phases. The rats in group M24M were deprived of food for 24 h before the expression phase test. Groups S0M, S0S, S24S, and S24M were treated with saline during the development phase. These groups were different because the rats in group S0M were treated with METH (1.0 mg/kg i.p.) prior to the expression phase test (day 8), the rats in group S0S were treated with saline prior to the expression phase, the rats in group S24S were deprived of food for 24 h before the expression phase test and treated with saline, and the rats in group S24M were deprived of food for 24 h before the expression phase test and treated with METH (1.0 mg/kg). Behavioral activity was recorded for 60 min in open-field cages, and blood glucose concentration was measured on days 1, 5 and 8 before the experimental session (Figure 1B shows the experimental design). Blood was collected from the tail vein, and glucose concentration was measured with a glucometer (Advantage Accu-Chek, Roche Diagnostics). The group names correspond to the treatment received during the experiment (M: methamphetamine; S: saline; 0 or 24 h of food deprivation).

**Figure 1:**
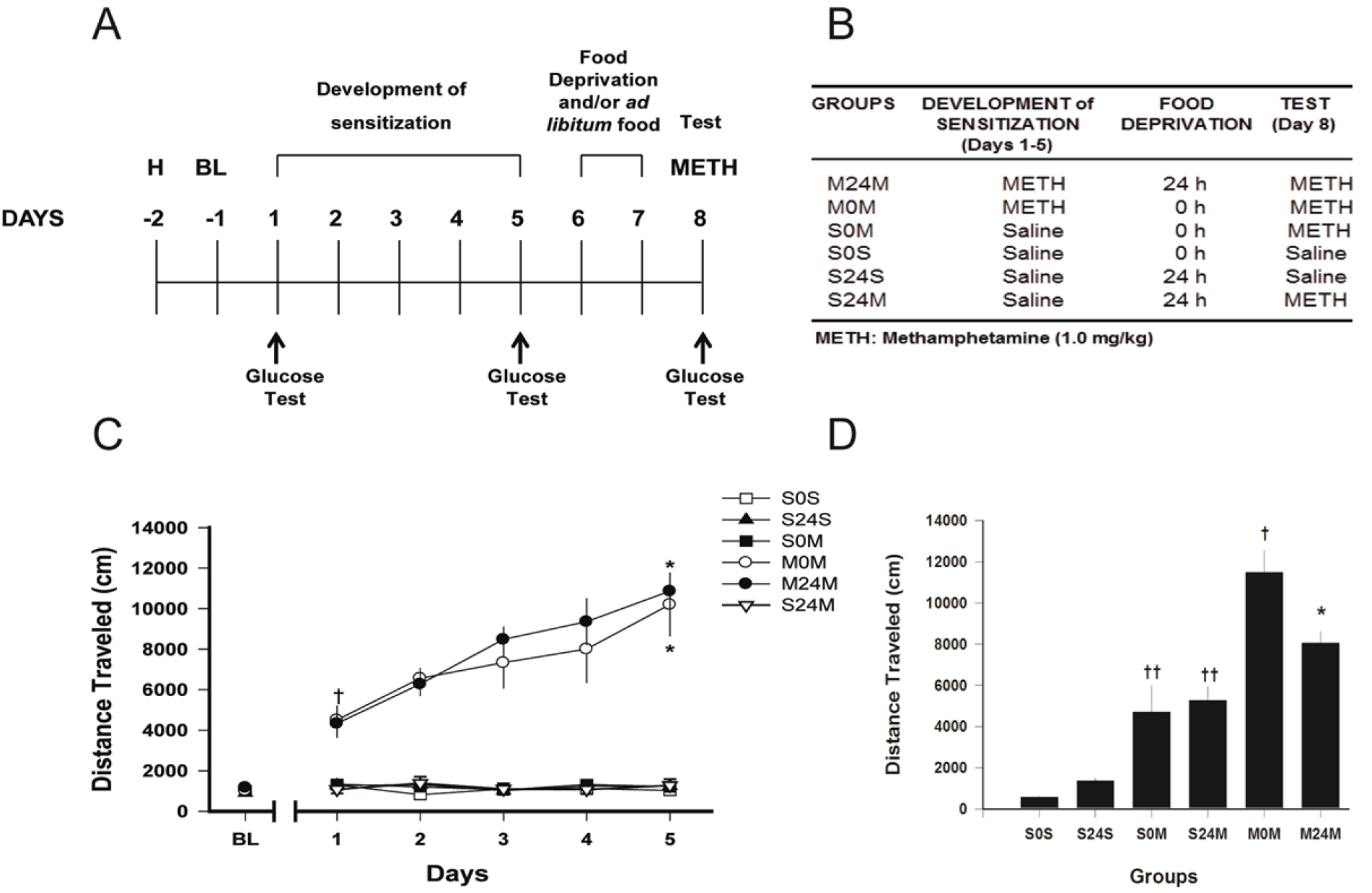
Effects of food deprivation on METH-induced locomotor sensitization. A. Schematic diagram illustrating the timeline for METH (1 mg/kg)-induced locomotor sensitization and the food deprivation schedule. Rats received repeated injections of METH or saline for 5 days. Some rats were deprived of food 24 h before the expression test (see Figure 1B). H, habituation; BL, baseline locomotor activity; METH, methamphetamine. B. Experimental design for the development and expression of METH-induced locomotor sensitization. C. Development of METH-induced locomotor sensitization in rats. METH (1.0 mg/kg) or saline was injected into animals daily for 5 days. Points are means ± SEM of 10 rats, asterisks (*) indicate significant differences between day 5 vs day 1, and a cross (†) indicates significant differences between the M0M, M24M groups and S0S S0M, S24S, and S24M groups on day 1 revealed by two-way ANOVA for repeated measures followed by Tukey’s post hoc test. D. Effects of food deprivation on the METH sensitization test. Rats were injected with saline or METH (1.0 mg/kg) on day 8. Bars represent the means ± SEM of 10 rats, an asterisk (*) indicates significant differences between the M24M and M0M groups, a cross (†) indicates significant differences between the M0M group and all other groups, and a double cross (††) indicates significant differences between the S0M and S24M groups and S0S and S24S groups based on one-way ANOVA followed by Tukey’s post hoc test.

### 2.5. Quantification of Insulin Levels by ELISA

Independent groups of rats were food deprived for 0 or 24 h, and their insulin levels were measured with a Rat Insulin ELISA Kit (Alpco Diagnostics, Windham, NH, USA) according to the manufacturer’s instructions. Blood samples were obtained from rats anesthetized with an i.p. injection of 2.5% avertin (Sigma-Aldrich, Mexico) before heart puncture, and then the samples were centrifuged to obtain blood serum. Briefly, 10 μl of each standard, control, and serum sample were placed on a microplate coated previously with a specific insulin monoclonal antibody, and then 75 μl of Working Strength Conjugate was added. The microplate was covered and incubated for 2 h at room temperature. After incubation, the microplate was washed 6 times with Working Strength Wash Buffer, 100 μl of tetramethylbenzidine substrate was added to each well, and the plate was incubated for 15 min at room temperature. Then, 100 μl of Stop Solution was added to each well. The absorbance of the enzymatic reaction was measured at 450 nm in an ELISA reader (Thermo Lab Systems Multiskan Ascent). All determinations were performed in duplicate. The intensity of the color generated was directly proportional to the amount of insulin in the sample.

### 2.6. Western Immunoblotting to Detect Phosphorylation of Akt at Ser473

Independent groups of rats were food deprived or not for 24 h, anesthetized with Halothane in a sealed induction chamber and sacrificed by decapitation. The brain was quickly removed and immersed in ice-cold (4°C), low-calcium ACSF containing 130 mM NaCl, 3 mM KCl, 5 mM MgCl2*5H2O, 26 mM NaHCO3, 1.25 NaH2PO4, 1 CaCl2, 10 glucose. Brain slices (500 μm) containing the nAcc were obtained using a vibratome (PELCO, 1000 Plus Sectioning System, Ted Pella Inc., Redding, CA, USA), and nAcc tissue was dissected and collected. nAcc tissues were homogenized in lysis buffer containing Tris-HCl 26 mM, 1% Triton x-100, glycerol 1.3 M, NaCl 130 mM and a protein phosphatase inhibitor cocktail Complete mini tab. The samples were collected and centrifuged for 5 min at 4500 rpm, and the supernatant was collected and stored at −70°C. Protein quantification was performed with the Bradford method, and 50 μg of protein was loaded in 10% polyacrylamide gels for electrophoresis. Proteins were transferred to PVDF transfer membranes and incubated with primary antibodies against pAKT serine 470 (Cell Signaling Technology) (1:1000) and β-tubulin (Cell Signaling Technology) (1:1000) for 12-20 h at 4°C and with secondary antibodies with horseradish peroxidase for 2 h at room temperature. Protein detection was performed using the chemiluminescence method with the aid of a C-Digit western blot scanner. Data were analyzed using Image Studio Digits 4.0 software. Immunoblotting experiments were performed in triplicate.

### 2.7. Data Analysis

The results of the investigated measure (distance traveled, cm) were expressed as the mean ± SEM. The data obtained during the development of locomotor sensitization were analyzed using two-way analysis of variance (ANOVA) for repeated measures, with day (1-5) as the repeated measure factor and groups as the between-subjects factor. The data obtained during the baseline locomotor activity and testing day (day 8) were analyzed through one-way ANOVA. One-way ANOVA was also used to assess the data obtained for blood glucose levels (on days 1, 5, and 8). When the ANOVA results were significant, Tukey’s test (p < 0.05) was used to perform a posteriori comparisons. An unpaired *t*-test was performed to analyze the effects of acute food deprivation on Akt activity in the rat nAcc and the effects of food deprivation on insulin levels.

## 3. Results

### 3.1 Repeated Administration of METH Produces Locomotor Sensitization

The data obtained from the baseline locomotor activity measurements were similar across all groups [F (5, 54) = 0.660, p = 0.655]. Repeated administration of METH resulted in the development of sensitization of locomotor activity in the M24M and M0M groups (Figure 1C), while repeated saline administration did not alter locomotor activity in the S0M, S0S, S24S and S24M groups. Two-way ANOVA for repeated measures indicated significant effects of group [F (5, 54) = 120.959, p = 0.0001], day [F (4, 216) = 9.855, p = 0.0001], and a group × day interaction [F (20, 216) = 4.358, p < 0.001]), and Tukey’s test revealed that the M24M and M0M groups were different from all of the other groups. Tukey’s test also revealed that the M24M and M0M groups were different from the S0M, S0S, S24S, and S24M groups on day 1, and also revealed significant differences between day 5 and day 1 on M24M and M0M groups. The results of food deprivation on the METH sensitization test on day 8 are shown in Figure 1D; food deprivation produced a reduction in METH-induced locomotor activity [F (5, 54) = 25.416 p = 0.0001]. Furthermore, Tukey’s test revealed that the M24M group was different from the M0M group and that the M0M group was different from the S0S, S24S, S0M, and S24M groups. Administration of an acute dose of METH induced a significant increase in locomotor activity in the rats in the S0M and S24M groups compared with that of the S0S and S24S groups (one-way ANOVA followed by Tukey’s post hoc test).

### 3.2. Food Deprivation Decreases Glucose Levels

Figure 2A shows the results of the glucose level measurements during days 1, 5, and 8. Glucose levels did not differ between groups on day 1 [F(5,54) = 0.551, p = 0.738] and day 5 [F(5,54) = 0.547, p = 0.740]. However, food deprivation reduced glucose levels on day 8 [F(5,54) = 37.842, p = 0.0001], and Tukey’s test revealed that regarding glucose levels, the S24S group was different from the S0S group, the M24M group was different from the M0M group, and the S24M group was different from the S0M group.

**Figure 2:**
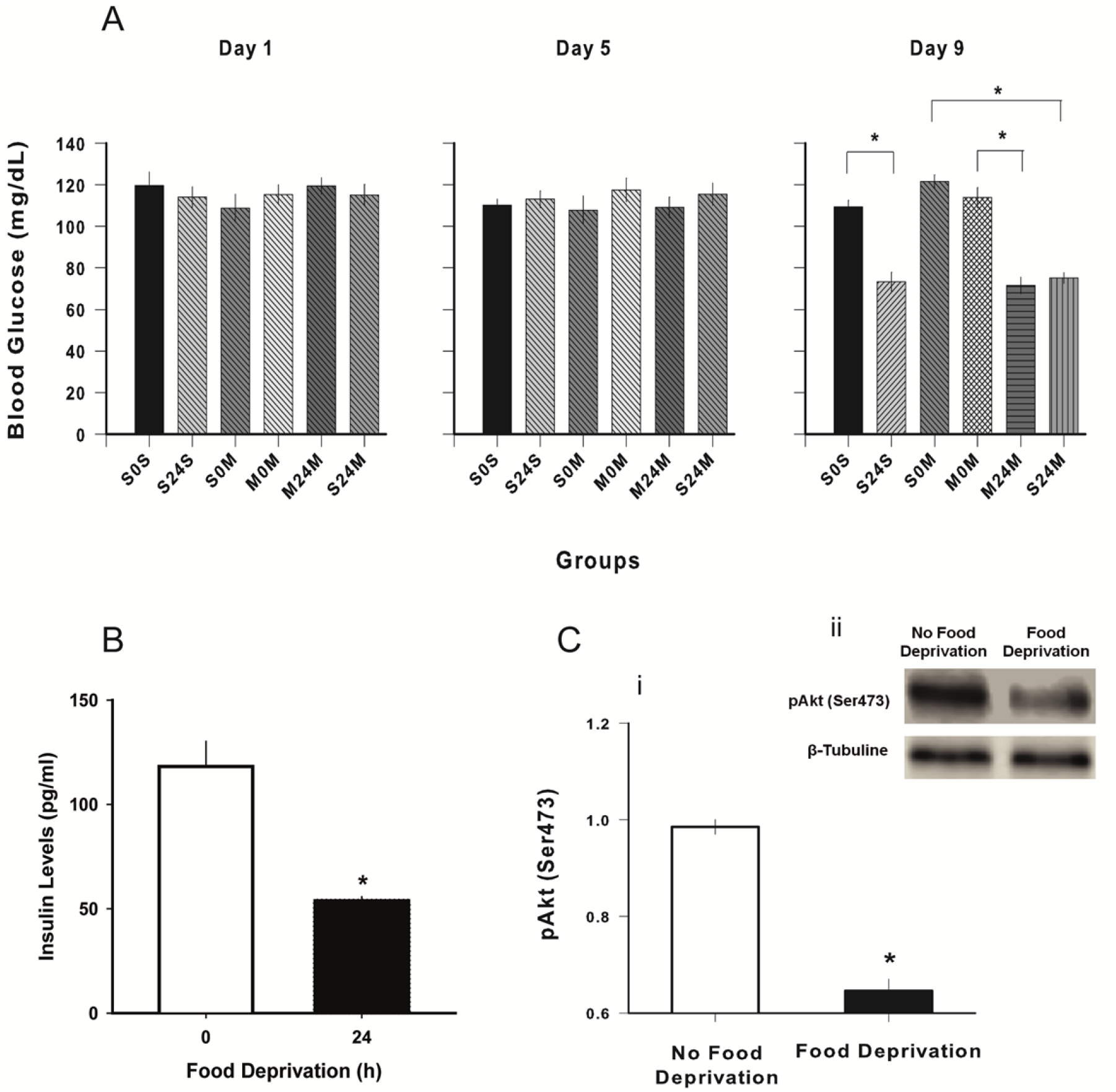
Effects of food deprivation on glucose, insulin, and Akt (Ser473) levels. A. Blood glucose levels in rats on days 1, 5 and 8. Bars represent the means ± SEM of 10 rats, and asterisks indicate significant differences between groups revealed by one-way ANOVA followed by Tukey’s post hoc comparisons. Blood glucose levels were measured with a glucometer (Roche: Accu-Check). A small drop of blood obtained by pricking the rat tail was placed on a glucometer. B. Insulin levels after 0 and 24 h of food deprivation. Bars represent the means ± SEM of 10 rats, and asterisks indicate significant differences compared to the 0 h of food deprivation revealed by Student’s *t*-test for independent samples. C. Food deprivation decreases pAkt (Ser473). i: Normalized data are expressed as the mean ± SEM. Asterisks indicate significant differences compared to non-food-deprived condition (Student’s *t*-test for independent samples [n = 3]). ii: qualitative representation of pAkt (Ser473) and β-Tubuline immunoreactive bands used for quantitative data analysis from nAcc.

### 3.4. Food Deprivation Decreases Insulin Levels

Figure 2B illustrates the effects of food deprivation on insulin levels. Food deprivation induced a significant reduction in insulin levels compared to nondeprived animals [t18 = 5,322, p = 0.0001].

### 3.5. Food Deprivation Decreases Phosphorylation of Akt

The phosphorylation levels of Akt at the serine 473 residue were considerably lower in the food-deprived group than in the control group (Figure 2C). Student’s *t*-test revealed that the food-deprived group was different from the non-food-deprived group [t4 = 11.952, p = 0.0001].

## 4. Discussion

The purpose of the present study was to examine the effects of acute food deprivation on METH-induced locomotor sensitization. We found that repeated METH administration increased locomotor activity and that acute food deprivation decreased METH-induced locomotor sensitization. We also observed that food deprivation reduced glucose and insulin levels. In addition, food deprivation also decreased phosphorylation of Akt at Ser473. The behavioral results described above are consistent with previous studies demonstrating that rats repeatedly injected with 1.0 mg/kg METH exhibited greater locomotor activity than saline-treated rats (Gu, Kim, Lamichhane, Hong, & Yun, 2019; Hall, Stanis, Marquez Avila, & Gulley, 2008; Kufahl et al., 2013; Wearne et al., 2015; Zhao et al., 2014). In the present study, it was also observed that acute food deprivation resulted in a decrease in METH-induced locomotor sensitization. Acute food restriction was scheduled for the 24 h prior to the expression test of METH-induced locomotor sensitization. These results are consistent with the few previous studies suggesting that food restriction plays an important role in regulating psychostimulant-induced behavioral effects. For example, AMPH (1.7-3.4 mg/kg but not 0.85 mg/kg)-induced conditioned place preference has been reported to be reduced by moderate food restriction, 15 g/day (Stuber, Evans, Higgins, Pu, & Figlewicz et al., 2002). In contrast to the present results and the findings mentioned above, some studies have reported that food restriction increased AMPH (1.7-3.4 mg/kg)-induced locomotor activity (Stuber et al., 2002). In addition, AMPH-induced locomotion has been reported to be unaffected by modest food restriction (15 g/day) (Sevak et al., 2008). The reason for this discrepancy may be related to acute food deprivation versus the food restriction regimen. While on a food restriction regimen, daily food intake is limited to a percentage of the food that is consumed by ad libitum-fed animals for several days, whereas in an acute food deprivation procedure, animals are maintained for several days with ad libitum food access and food deprived for a certain time, for example, 24 h, before behavioral and/or biochemical tests. This discrepancy seems to suggest that the acute food deprivation procedure reduces the expression of psychostimulant-induced locomotor sensitization and that the food restriction regimen increases rather than decreases the induction of psychostimulant-induced locomotor sensitization (see below for explanation).

The mechanism underlying the observed effects of food deprivation on METH-induced locomotor sensitization may involve insulin signaling modulation of psychostimulant-induced behavioral effects. Several lines of evidence support this notion. First, AMPH-like drugs are well-known DAT substrates that competitively inhibit DA uptake by increasing extracellular DA concentration (Khoshbouei, Wang, Lechleiter, Javitch, & Galli et al., 2003; Proebstl et al., 2019) and require the DAT for their action (Giros, Jaber, Jones, Wightman, & Caron, 1996; Proebstl et al., 2019). The DAT is a plasma membrane transport protein that belongs to the Na+/Cl-dependent transporter gene family and is responsible for regulating the concentration of extracellular DA (Giros & Caron, 1993; Vaughan & Foster, 2013). Second, AMPH increases DA release, depleting vesicular DA stores and promoting DA release through reversing the DAT function (Fleckenstein, Volz, Riddle, Gibb, & Hanson, 2007; Karam & Javitch, 2018; Sulzer, 2011). AMPH-like drugs not only inhibit DA uptake and increase DA efflux mediated through the DAT but also decrease DAT surface expression (Fleckenstein, Metzger, Wilkins, Gibb, & Hanson,1997; Kahlig et al., 2006; Karam & Javitch, 2018; Saunders et al., 2000). These effects of AMPH-like drugs produce an increase in DA neurotransmission (Wheeler et al., 2015). Third, several signal transduction pathways have been reported to be able to modulate DAT trafficking and activity, including insulin signaling cascades (Daws et al., 2011; Fiory et al, 2019). Insulin can cross the bloodbrain barrier (Rhea, Rask-Madsen, & Banks, 2018; Schulingkamp, Pagano, Hung, & Raffa, 2000) by a saturable mechanism (see Banks, Owen, & Erickson, 2012) and acts on insulin receptors in the limbic areas of the brain such as the striatum (Fiory et al, 2019; Schulingkamp et al., 2000), a brain region in which the DAT is highly expressed (Figlewicz, Evans, Murphy, Hoen, & Baskin, 2003). Insulin depletion by streptozotocin, a toxin that selectively destroys the b-cells of the pancreas (Dulin & Soret, 1977; Zhang et al., 2018), is reported to reduce DAT activity (Owens et al., 2005; Sevak et al., 2007) and surface expression (Fiory et al, 2019; Williams et al., 2007) and reduce AMPH self-administration behavior (Galici et al., 2003; Williams et al., 2007). Like insulin depletion by streptozotocin, food restriction also reduces blood insulin levels (Carr, 1996; Stouffer et al., 2015) and DAT activity (Patterson et al., 1998; Sevak et al., 2008; Jones, Woods, Zhen, Antonio, Carr, & Reith, 2017; Zhen, Reith, & Carr, 2006). In the present experiment, food deprivation for 24 h also reduced glucose and insulin levels (see Figure 2A and B). These results are in agreement with experiments that reported that blood glucose and insulin levels displayed similar patterns of change in rats that were subjected to different food restriction treatments (Mamczarz et al., 2005; Marinković et al., 2007).

Insulin pathways have been suggested to play an important role in regulating DAT function and reward pathways that are involved in the behavioral actions of AMPH-like drugs (Daws et al., 2011; Fiory et al, 2019). Activation of insulin signaling pathways, including Akt, a protein kinase immediately downstream of PI3K, is essential for regulating DA clearance and surface expression of the DAT (Fiory et al, 2019; Garcia et al., 2005). Inhibition of Akt via ML9 reduced surface expression of the DAT and the ability of insulin to modulate AMPH-induced DAT surface redistribution (Garcia et al., 2005). This evidence suggests an important role for insulin and its signaling pathways in DAT function and the ability of AMPH-like drugs to increase DA neurotransmission. In the present experiment, we reported that food deprivation also decreased Akt phosphorylation at Ser473 and that this effect could be related to a decrease in METH-induced locomotor sensitization. In line with these results, it has been reported that the integrity of PI3K, a protein kinase activated by insulin signaling, is required for the expression, but not the induction, of locomotor sensitization to cocaine (Izzo et al., 2002) and cocaine-seeking behavior (Szumlinski, 2019)

In conclusion, the results of the present study showed that food deprivation for 24 h reduced METH-induced locomotor sensitization. Furthermore, food deprivation also reduced levels of blood glucose, insulin and phosphorylation of Akt at Ser473. These data provide further evidence that insulin and the protein kinase Akt may modulate AMPH-like drug-induced behaviors, indicating the potential utility of insulin signaling pathways to treat psychostimulant addiction.

## Authors’ contribution

F.M. Conceived and designed the experiments. J.C.J. Performed the behavioral experiments. J.A.M-B. Performed Immunoblotting experiments. M.B-D. Performed quantification of insulin level experiments. F.M. and J.C.J. Analyzed data and wrote the manuscript, with contributions of all authors.

## Acknowledgments

This study was supported by grant IN307414 from PAPIIT-UNAM (México).

